# Domain-specific functions of LRIT3 in synaptic assembly and retinal signal transmission

**DOI:** 10.1101/2025.09.21.677564

**Authors:** Nazarul Hasan, Ronald G. Gregg

## Abstract

LRIT3 is a leucine-rich repeat (LRR) protein that is expressed in the retina, and its absence causes complete congenital stationary night blindness (cCSNB), a genetically diverse disorder characterized by impaired low-light vision, myopia, and nystagmus. LRIT3 is expressed in rod and cone photoreceptors, and it trans-synaptically organizes the assembly of the glutamate signaling complex, the signalplex, on depolarizing bipolar cells (DBCs). LRIT3 is a single-pass membrane protein with extracellular LRR, IG, and FN3 domains. The mechanism by which LRIT3 controls postsynaptic receptor organization remains unknown. We address this by using rAAV to express deletion constructs in LRIT3 knockout retinas and examining LRIT3 trafficking, as well as the structural and functional recovery of the signalplex in DBCs. We show the LRR domain is required for trafficking LRIT3 to the synapse in cones, but not rod photoreceptors, although it is needed for reassembly and function of the rod BC signalplex. Neither the IG nor the FN3 domain is needed for synaptic localization of LRIT3. However, the IG domain is required for the localization of TRPM1 to the signalplex and thus function. The FN3 domain is not necessary for either DBC signalplex assembly or function. Our data demonstrates that the LRR and IG domains of LRIT3 are crucial for TRPM1 localization and retinal function, with the LRR domain playing a key role in the differential function of LRIT3 at rod and cone synapses. Notably, our results show that restoring Nyctalopin localization to the DBC signalplex alone is insufficient to restore TRPM1 expression. Based on our findings, we propose a model in which the LRR domain trans-synaptically binds with Nyctalopin, while the IG domain interacts with TRPM1.

## Introduction

Vision begins in the retina with rod and cone photoreceptor cells, which are specialized for detecting light. When light levels increase, these photoreceptors hyperpolarize and reduce their release of glutamate in the outer plexiform layer (OPL). This change is detected by second order neurons, bipolar (BC) and horizontal cells. Of the 14 bipolar types, 5 hyperpolarize (HBCs) in response to a light increment and 9 depolarize (DBCs). Rod photoreceptors primarily connect to a single type of DBC, while cones connect to 8 types of DBCs and 5 types of HBCs. DBCs detect changes in glutamate release through the metabotropic glutamate receptor 6 (mGluR6), which modulates the non-specific cation channel TRPM1 (1, 2). Cones connect to two types of bipolar cells: DBCs, which also use mGluR6, and HBCs that use AMPA/kainate ionotropic glutamate receptors (3–6). Rod bipolar cells and horizontal cells form a single invaginating synapse near a ribbon structure within the rod photoreceptor spherule. Similarly, cone pedicles have multiple invaginating synapses involving various classes of cone DBCs and horizontal cell dendrites, all positioned near a ribbon. In contrast, HBCs form flat contacts on the base of the cone pedicle. This intricate synaptic organization allows the retina to process and transmit visual information efficiently, enabling us to see in both dim and bright light conditions.

Anatomically, invaginating synapses of photoreceptors are intricate structures involving multiple pre- and postsynaptic proteins, including several adhesion molecules that maintain synapse stability and functionality (7–11). Despite knowing many of the involved components, the molecular mechanisms underlying synapse formation and receptor complex assembly remain poorly understood. Mutations in *GRM6*, *TRPM1*, *Nyctalopin*, *GPR179*, and *LRIT3* have been identified as disruptors of ON pathway signaling in humans, leading to complete congenital stationary night blindness (cCSNB, see review by Zhang et al. (12). In mice, these have all been shown to be part of a large signalplex located on the tips of the DBC dendrites.

LRIT3 is a leucine-rich repeat (LRR) protein that is involved in transsynaptic organization of the DBC signalplex, is a single-pass membrane protein that contains an extracellular LRR, IG and FN3 domains. Three other LRR-containing proteins, Nyctalopin, ELFN1, and ELFN2, also are involved in synaptic protein assembly and/or synaptic transduction in the OPL. LRIT3 is expressed exclusively at rod and cone synapses with DBCs (8, 9, 13). Functionally, the loss of LRIT3 results in a no b-wave phenotype in mice (8, 9). In the absence of LRIT3, there are differential effects on rod and cone DBC signalplexes. In rods, Nyctalopin and TRPM1 fail to localize at rod BC dendrites, and in cones the entire postsynaptic signalplex, including mGluR6, GPR179, Nyctalopin, TRPM1, and RGS7/11 is missing (9, 14, 15). Given these differences between rod and cones synapses in *Lrit3^-/-^*retinas, we hypothesized that the extracellular domains of LRIT3 contribute to its differential functions, with each domain playing a specific role in synaptic protein assembly.

We now demonstrate that the LRIT3 IG domain is crucial for the localization of TRPM1 at both rod and cone synapses, and the LRR domain is essential for the localization of Nyctalopin and TRPM1 at rod synapses and LRIT3 trafficking to cone synaptic pedicles.

## Results

Mutations in the *LRIT3* gene have been identified as a cause of complete congenital stationary night blindness in humans (16). The expression of full-length LRIT3 in cones or rods of *Lrit3^-/-^* mice restores function (9, 13, 17). Because LRIT3 has differential effects on rod and cone synapses (8, 9) and has three predicted functional domains, LRR, IGc2 (IG), and FN3, we asked whether these specific domains have distinct functions. To explore this, we created deletion constructs lacking the LRR, IG, or FN3 domains, as well as both IG and FN3. (Fig 1A). Because the LRIT3 antibody was generated against the LRR domain, in ΔLRR constructs, we inserted a Myc tag between amino acids 19 and 20 of LRIT3. To limit expression to either rods or cones, we used the RHO promoter (18) or the GNAT2 promoter (19), respectively. Figure 1B shows the general design of the AAV constructs packaged into rAAV8 capsids. The rAAVs were then injected into the subretinal space of *Lrit3^-/-^* mice on postnatal day ∼35 (P35), and protein trafficking and function were assessed at ∼P70, 5 weeks post-injection (Fig. 1C).

**Figure 1.**
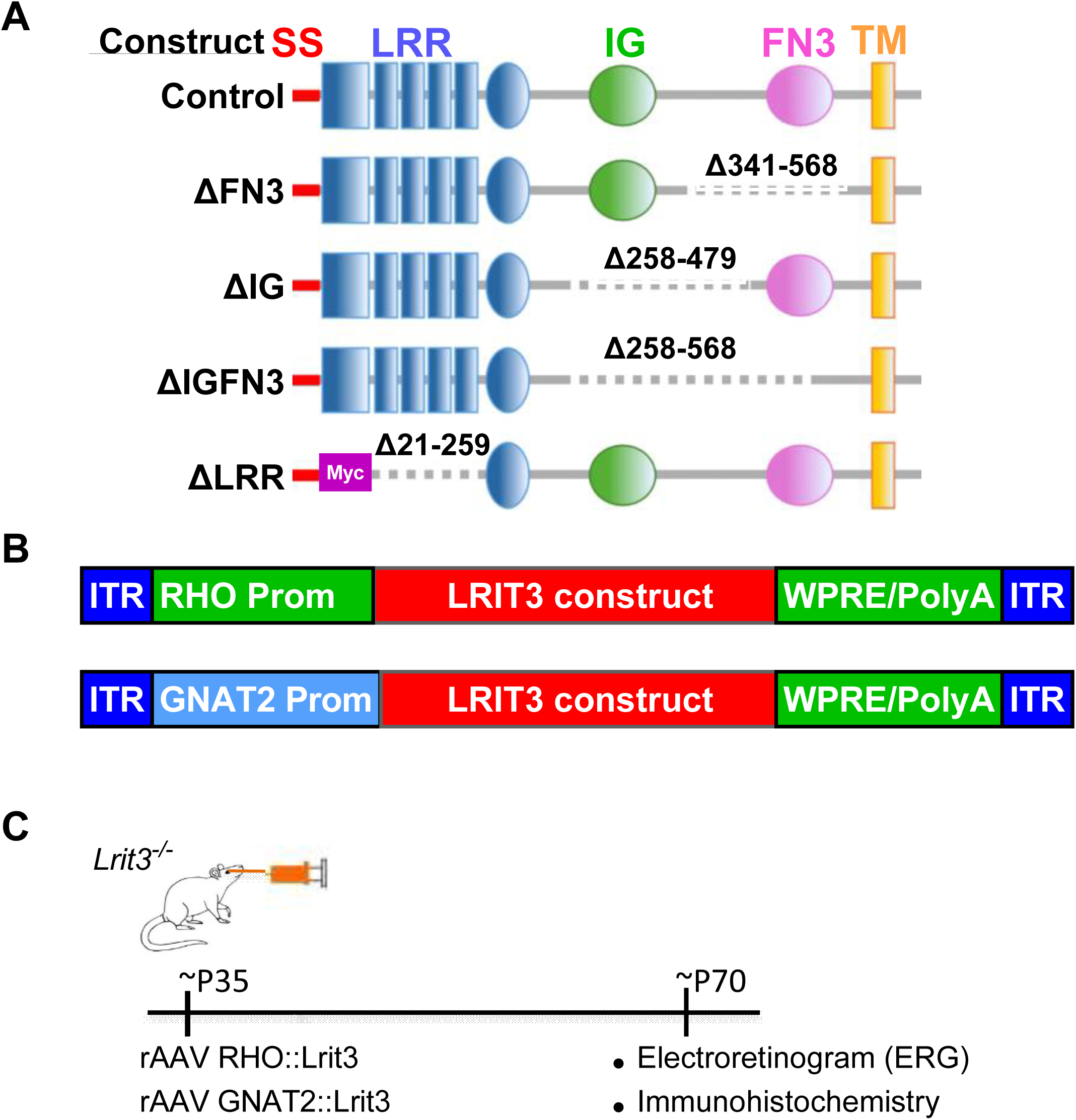
Experimental design and vector constructs for LRIT3 expression in mouse retina. (A) Structure-based design of LRIT3 protein constructs with systematic deletion of extracellular domains. Wildtype LRIT3 and three truncated variants with selective removal of fibronectin type III (FN3), immunoglobulin-like (IG), and leucine-rich repeat (LRR) domains, respectively. (B) Schematic of the two recombinant adeno-associated virus (rAAV) vectors containing rhodopsin (RHO) and GNAT2 promoter for targeting rod and cone photoreceptors, respectively. (C) Experimental workflow: Adult *Lrit3*^-/-^ mice received subretinal rAAV injections. After 5 weeks, retinal function was assessed by electroretinography (ERG) and protein expression was evaluated by immunohistochemistry.

### LRIT3 domains play differential roles in trafficking in rods and cones

The expression of deletion constructs used to probe the function of the various protein domains in LRIT3 may lead to abnormalities in protein trafficking. To address this, we used immunohistochemistry (IHC) to determine if trafficking was normal and whether other components known to be absent in *Lrit3^-/-^* mice (9, 14, 17) were re-localized to the rod and cone DBC signalplexes. We immunostained transverse sections of retinas injected with rAAVs expressing mutant forms of LRIT3 in rods or cones, and used wholemount retinas to quantify the infection frequency and trafficking. We considered colocalization of LRIT3 with mGluR6 in rods (Fig. 2) and with cone arrestin (mCAR) in cones (Fig. 3), to assess trafficking. Example images for all constructs are shown for a small section of a wholemount retina for each construct (Fig. 2,3). Colocalization from 5 regions of each retina/construct in 3 separate retinas was imaged, and the fraction of rods and cones infected was quantified (Fig. 2B, 3B). The images shown are representative of at least three retinas analyzed for all conditions. To further establish co-localization we used IHC to triple label retinas for LRIT3, Nyctalopin (Nyx-EYFP) and TRPM1 for rod (Fig. 4) and cone (Fig. 5) specific constructs. To establish co-localization, we created plots of puncta intensity for each of the three antibodies (graphs to the right of images in Fig. 4,5).

**Figure 2.**
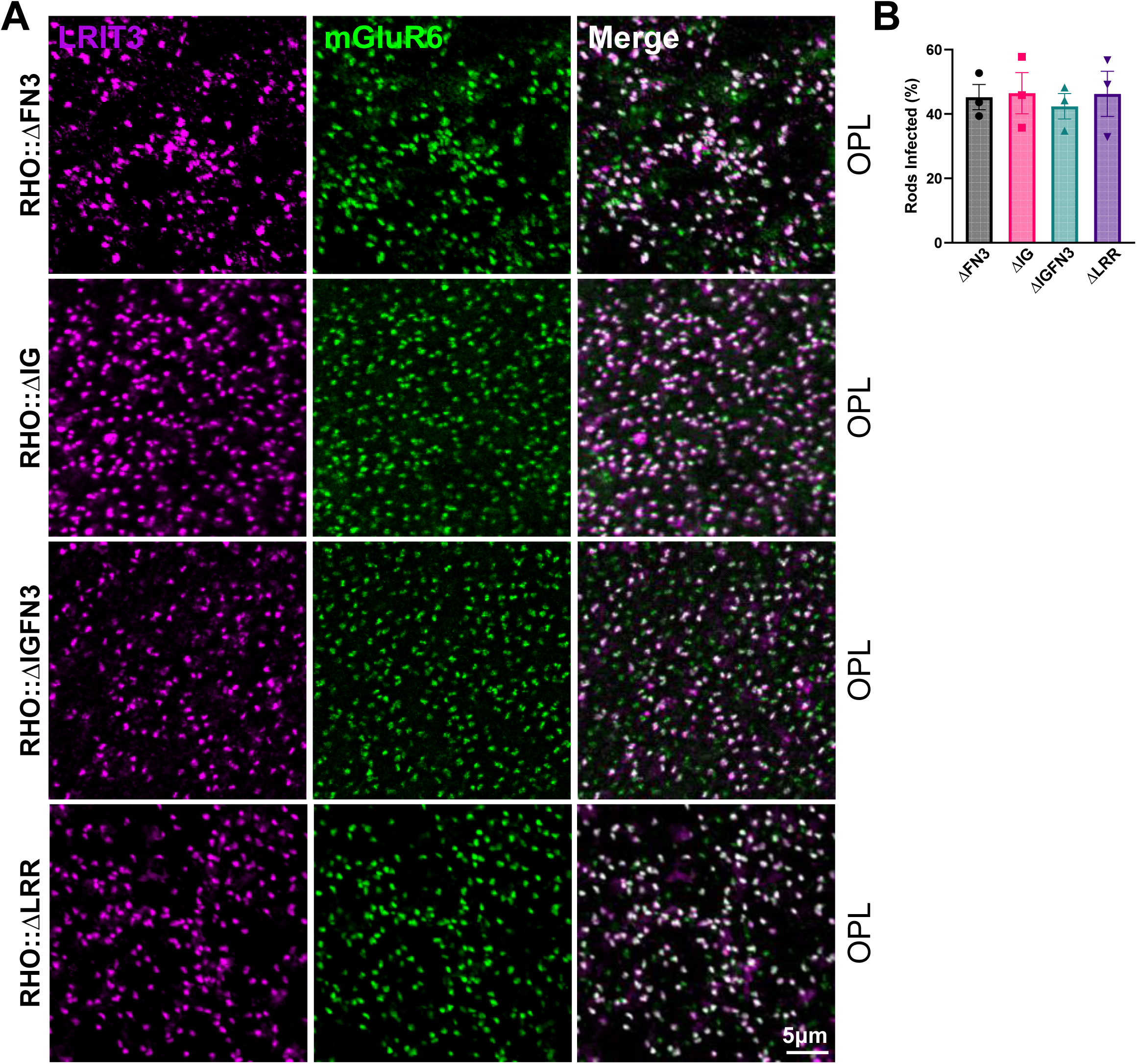
LRIT3 deletion constructs successfully traffic and localize at rod synapses. (A) Representative confocal images of wholemount retinas from *Lrit3^-/-^* mice treated with rAAV8 RHO::ΔFN3, rAAV8 RHO::ΔIG, rAAV8 RHO::ΔIGFN3, and rAAV8 RHO::ΔLRR. Images were captured at the outer plexiform layer (OPL) after staining with antibodies to LRIT3 (magenta) and mGluR6 (green). Scale bar = 5 μm. (B) Quantitative analysis of LRIT3 deletion construct expression at rod synapses. Bar graph shows the percentage of rod synapses positive for each deletion construct in *Lrit3^-/-^* retinas (n = 3 eyes from 3 mice per group).

**Figure 3.**
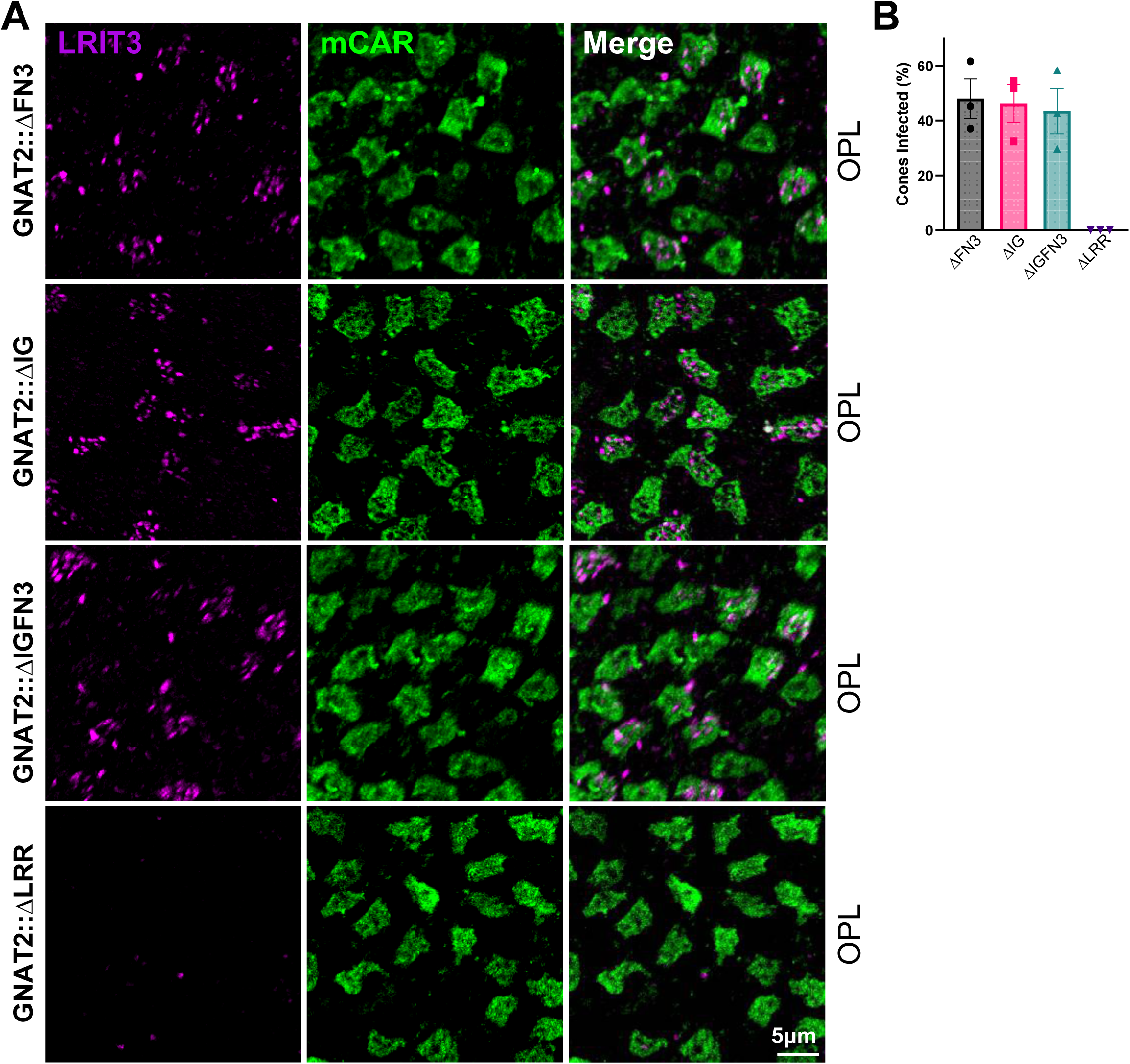
LRR domain is critical for LRIT3 localization at cone synapses. (A) Representative confocal images of wholemount retinas from *Lrit3*^-/-^ mice treated with rAAV8 vectors expressing GNAT2-driven LRIT3 deletion constructs (ΔFN3, ΔIG, ΔIGFN3, and ΔLRR). Images were taken at the OPL level following immunostaining using antibodies to LRIT3 (magenta) and mCAR (green). Scale bar = 5 μm. (B) Quantitative analysis of LRIT3 deletion constructs expression at cone synapses. Bar graph displays the percentage of cone synapses positive for each deletion construct in *Lrit3*^-/-^ retinas following treatmentt (n = 3 eyes from 3 mice per group). OPL = outer plexiform layer; mCAR = mouse cone arrestin.

**Figure 4.**
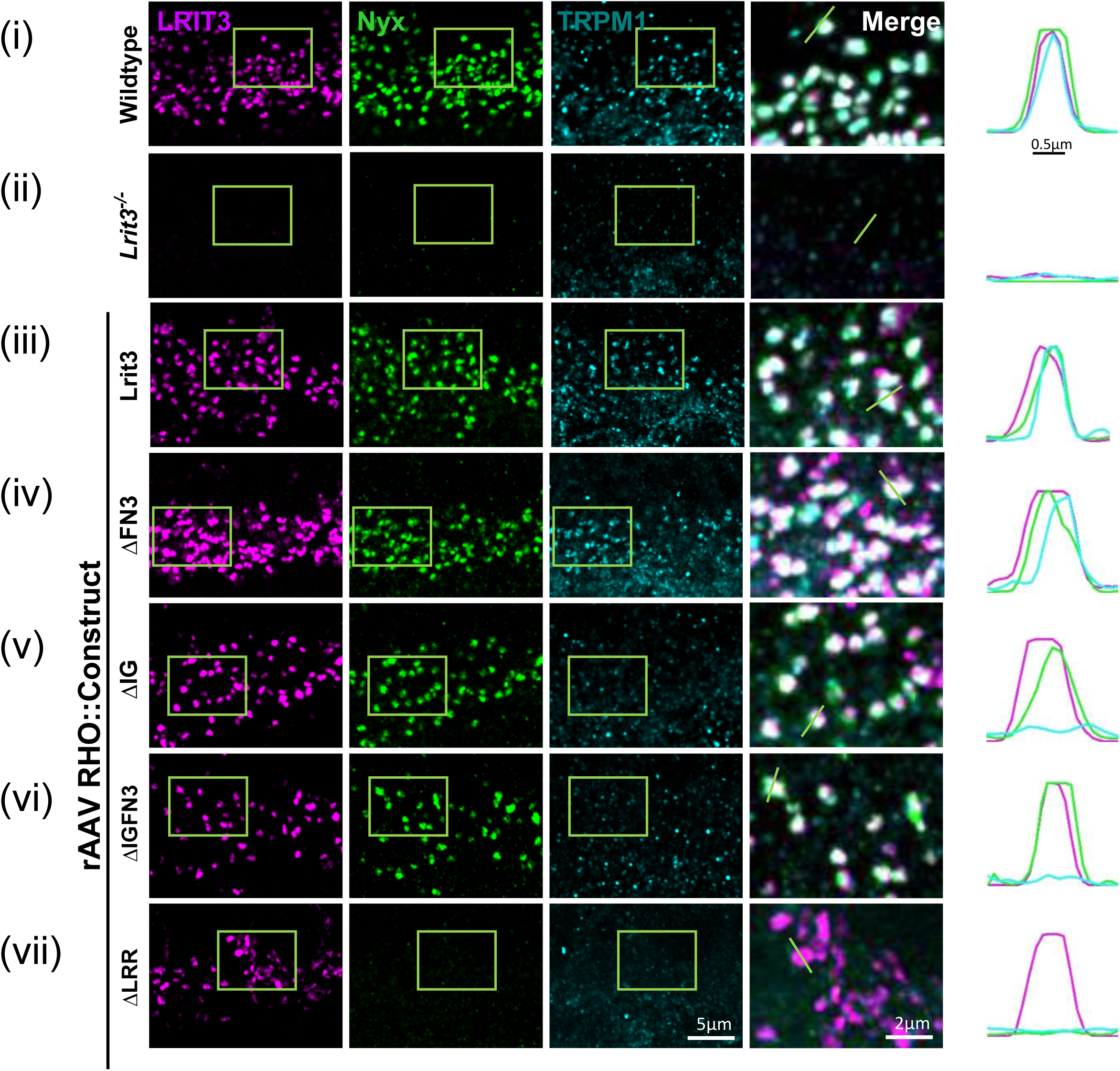
LRR and IG domains of LRIT3 are essential for TRPM1 localization at rod bipolar cell dendrites. (A) Representative confocal images of transverse sections of the outer plexiform layer from (i) Wildtype mice, (ii) *Lrit3^-/-^*, and (iii-vii) *Lrit3^-/-^* retinas treated with rAAV8 vectors expressing RHO-driven constructs: full-length Lrit3, ΔFN3, ΔIG, ΔIGFN3, and ΔLRR, respectively. Sections were immunostained using antibodies to LRIT3 (magenta), GFP-Nyctalopin (Nyx, green), and TRPM1 (cyan). Scale bar = 5μm. Boxed regions are shown as enlarged merged images from each condition. Scale bar = 2μm. Images are representative of n = 4 retinas per group. (B) Intensity profile plots (right panel) demonstrate the spatial relationship and colocalization of Nyctalopin, TRPM1. and LRIT3 puncta, determined by quantitative analysis of fluorescence intensity along the indicated green line in the corresponding merged images.

**Figure 5.**
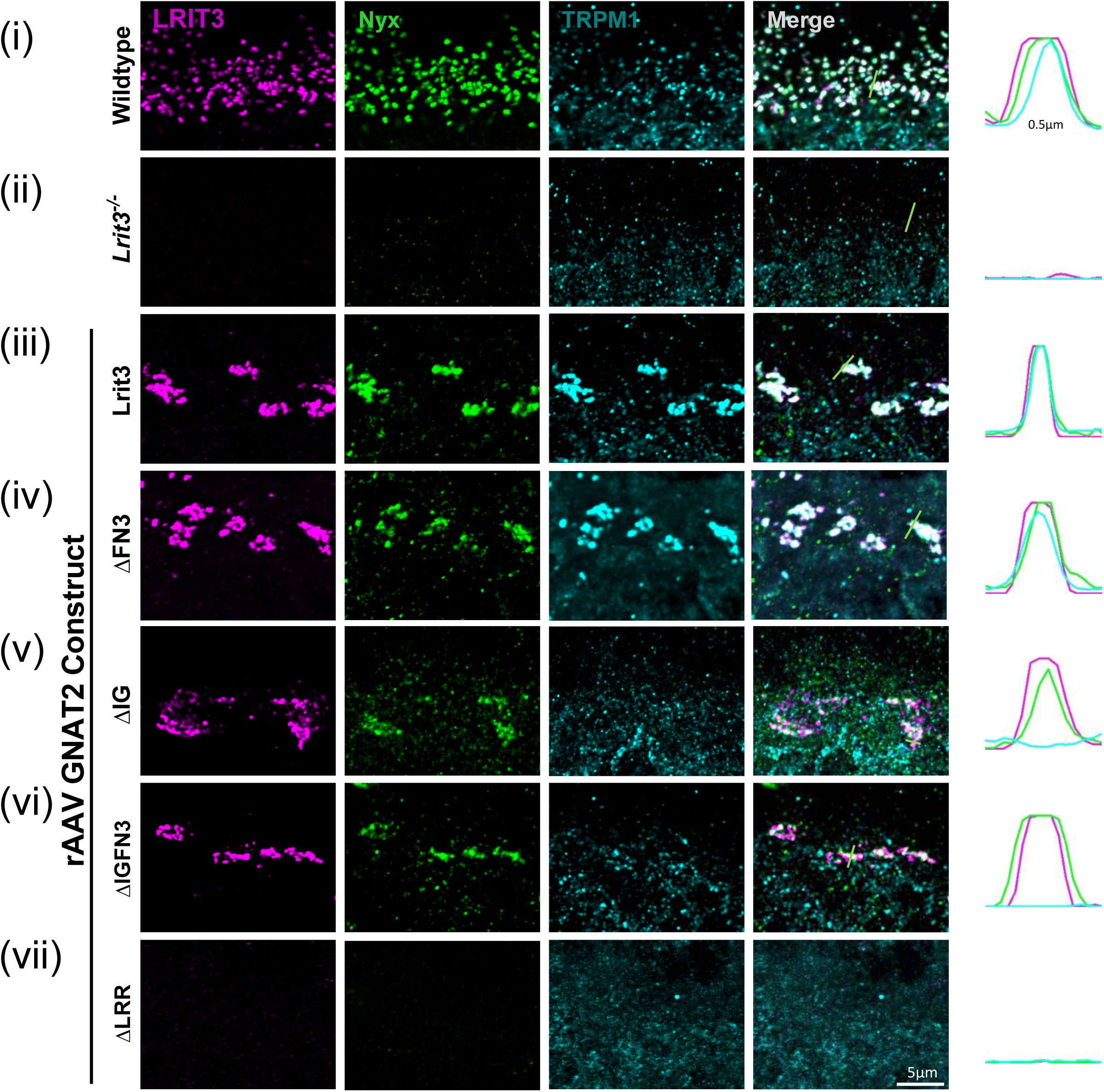
LRR domain of LRIT3 is essential for localization of LRIT3, nyctalopin, and TRPM1 at cone synapses. (A) Representative confocal images of transverse retinal sections of the outer plexiform layer from (i) Wildtype mice, (ii) *Lrit3*^-/-^, and (iii-vii) *Lrit3*^-/-^ retinas following subretinal delivery of rAAV8 vectors containing GNAT2 promoter-driven constructs: full-length Lrit3, ΔFN3, ΔIG, ΔIGFN3, and ΔLRR deletion mutants. Triple immunostaining for LRIT3 (magenta), GFP-Nyctalopin (Nyx, green), and TRPM1 (cyan). Scale bar = 5μm. Images are representative of four independent retinas per experimental group. (B) Quantitative intensity profile analysis shows spatial distribution and colocalization of Nyctalopin, TRPM1, and LRIT3 at cone synaptic terminals, measured along the designated green line in merged images.

When full-length LRIT3 or any of the deletion constructs were expressed in rods, they all trafficked as is evident from the punctate labelling seen for LRIT3 (Fig. 4). Except for the construct lacking the LRR domain (ΔLRR, Fig 4vii), Nyctalopin expression was also restored normally as indicated in the higher magnification merged images and colocalization plots for the ΔFN3, ΔIG, and ΔIGFN3 constructs. Thus, the LRR domain is essential for the correct localization of Nyctalopin. For TRPM1, the IG and LRR domains are required for proper localization, because it is not localized correctly when either of these domains is deleted.

Notably, for the ΔIG and ΔIGFN3 constructs, Nyctalopin is localized correctly, but TRPM1 is not, indicating our earlier hypothesis that Nyctalopin was sufficient to localize TRPM1 was incorrect (7). Rather, Nyctalopin and the LRIT3 IG and LRR domains are required for the localization of TRPM1 in rod BCs.

To examine the function of LRIT3 domains in cones, we treated *Lrit^3-/-^*retinas with rAAV using the GNAT2 promoter to drive expression of deletion constructs. We then immunostained for Nyctalopin (Nyx-EYFP), and TRPM1 after subretinal injection (Fig. 5). These images are representative of at least four separate retinas. The constructs lacking the IG and/or the FN3 domain trafficked and localized correctly, but unlike in rods, the ΔLRR domain construct did not traffic to the correct location on cone pedicles, so its function in assembling the cone DBC signalplex cannot be determined. Like the situation in rods, the IG and/or the FN3 domain was not needed for localization of ΔLRIT3s to the cone pedicle (Fig. 5iv-vi), and the FN3 domain is not required for Nyctalopin localization, but the LRR domain is. For TRPM1 localization, the IG domain was needed, based on a failure of the ΔIG (Fig. 5v), and the ΔIGFN3 (Fig. 5vi), constructs to localize TRPM1. Like the situation at rod pedicles, Nyctalopin localization is insufficient to restore TRPM1 to the signalplex of cone DBCs (Fig. 5iv,v).

In *Lrit3^-/-^* mice, mGluR6 and GPR179 are also missing from the dendrites of cone DBCs, but present on the rod BC dendrites. Therefore, we examined their localization after treatment of *Lrit3^-/-^* mice with cone specific GNAT2::ΔFN3, ΔIG and ΔIGFN3 constructs. In all cases, mGluR6 (Fig. 6) and GPR179 (Fig. 7) were restored to the cone DBC dendrites. Note that for both mGluR6 and GPR179, the green puncta in the merged image represent the rod BC dendrites, which does not require LRIT3 for its localization in the *Lrit3^-/-^* retinas. Whether GPR179 and mGluR6 require the LRR domain for localization in cone DBCs is currently unknown, due to the failure of the ΔLRR construct to traffic to cone synapses (Fig. 5vii).

**Figure 6.**
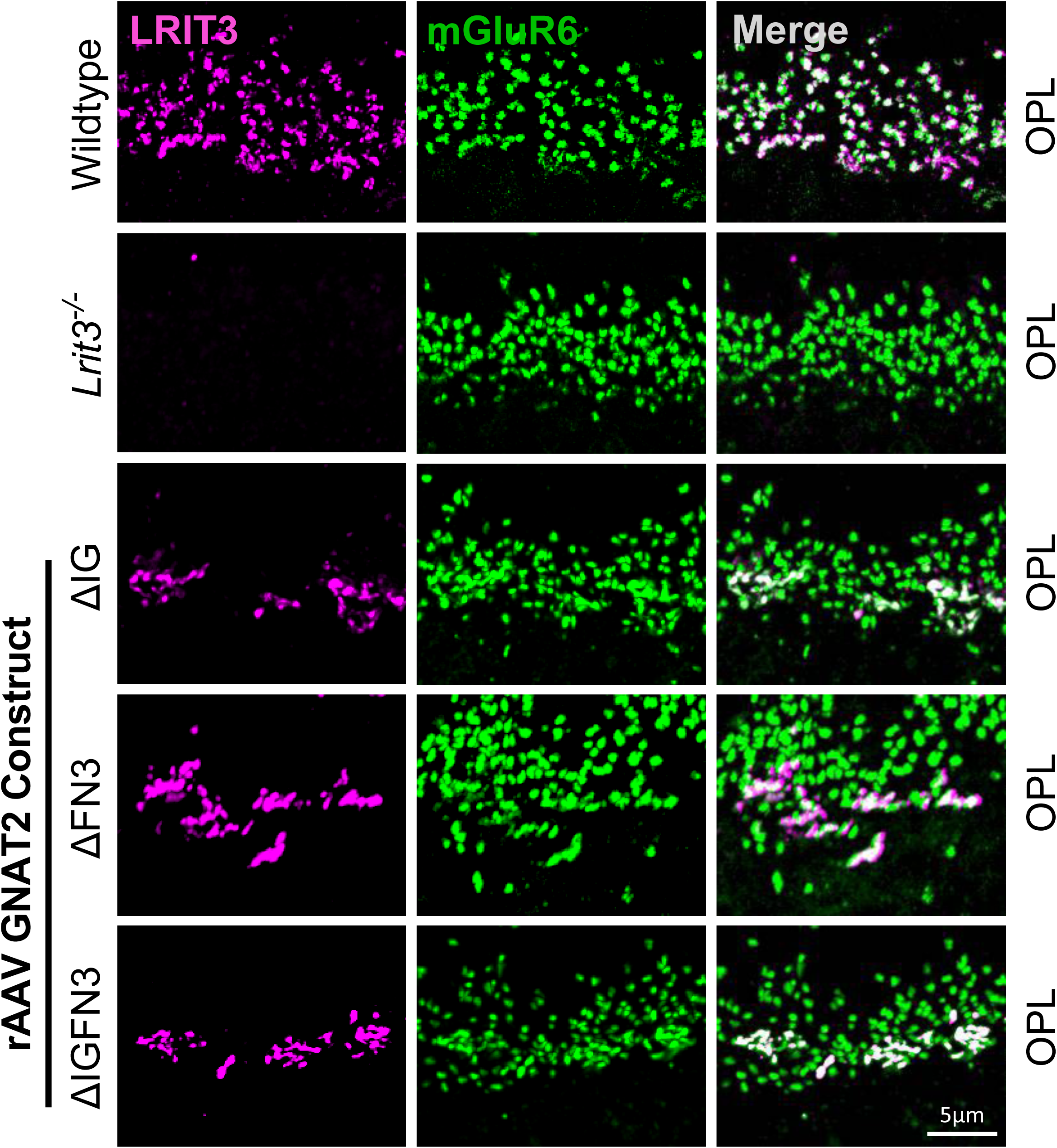
mGluR6 localizes at cone bipolar cell dendrites in the absence of IG and FN3 LRIT3 domains. Representative confocal images of transverse retinal sections from the outer plexiform layer of Wildtype mice, *Lrit3*^-/-^, and *Lrit3*^-/-^ retinas treated with subretinal injections of rAAV8 vectors expressing GNAT2-driven LRIT3 deletion constructs: ΔIG, ΔFN3, and ΔIGFN3, respectively. Sections were immunostained with antibodies against LRIT3 (magenta) and mGluR6 (green). Scale bar = 5μm. Images are representative of n = 4 retinas per group.

**Figure 7.**
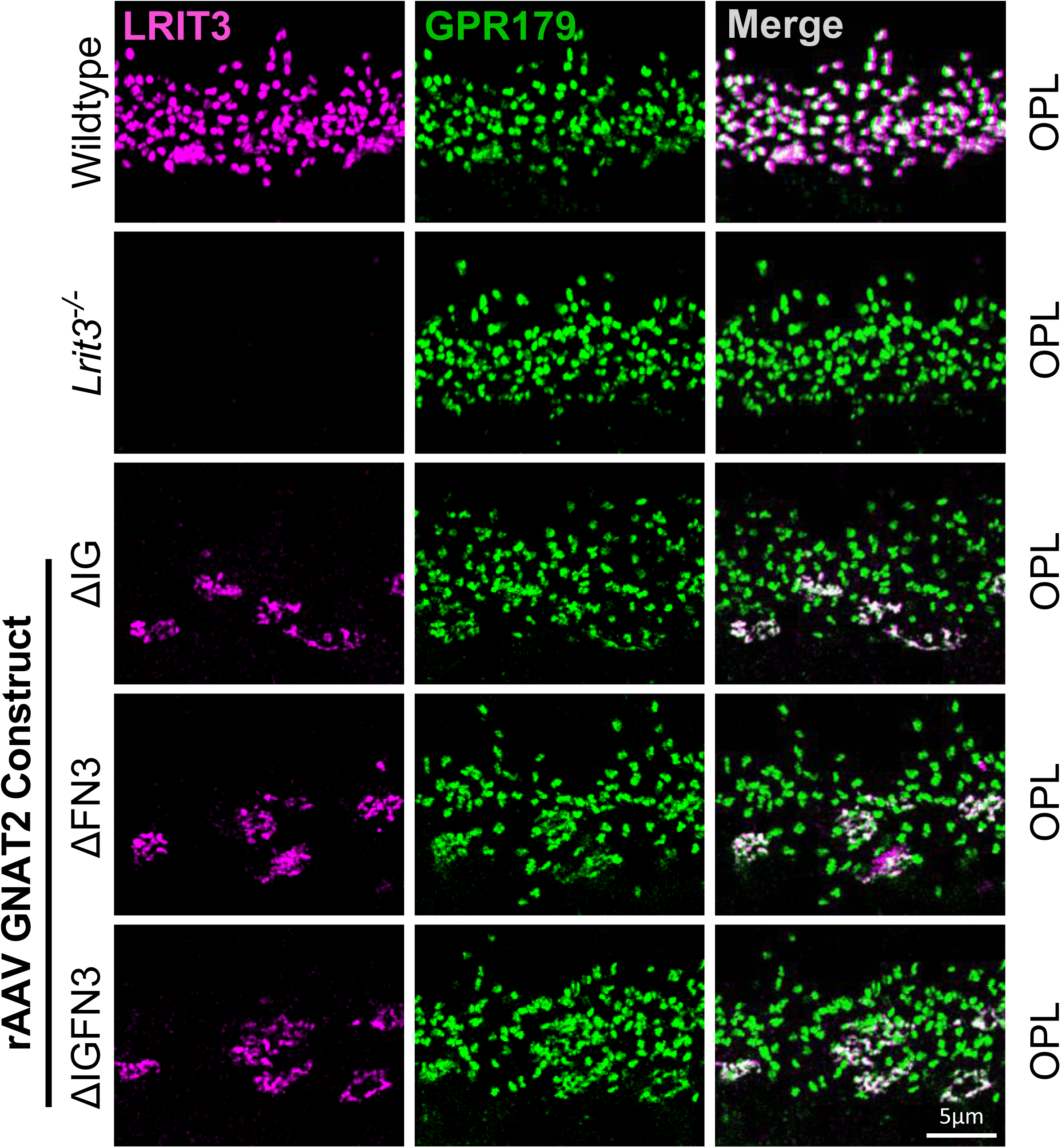
GPR179 localizes at cone bipolar cell dendrites in the absence of IG and FN3 LRIT3 domains. Representative confocal images of transverse retinal sections from the outer plexiform layer of Wildtype mice, *Lrit3*^-/-^, and *Lrit3*^-/-^ retinas treated with subretinal injections of rAAV8 vectors carrying GNAT2-controlled LRIT3 constructs lacking specific domains: ΔIG, ΔFN3, and ΔIGFN3. Sections were immunostained with antibodies against LRIT3 (magenta) and GPR179 (green). Scale bar = 5μm. Images are representative of n = 4 retinas per group Scale bar = 5μm.

### The LRIT3 FN3 domain is not required for DBC function

Signaling between photoreceptors and DBCs can be assessed using the electroretinogram (ERG). The a-wave represents photoreceptor function, and the b-wave the response of DBCs. When measured under scotopic or photopic conditions, it provides functional information about the DBC signalplex in rod and cone DBCs, respectively. The IHC above showed that expression of only the full-length LRIT3 and the ΔFN3 construct restored TRPM1 expression to the DBC dendrites of rod BCs and cone BCs. This would predict that only these two treatments would restore function. To address this question, treated animals were assessed with the electroretinogram, which measures the depolarization of DBCs, after overnight dark adaptation. We measured the scotopic ERG b-wave to assess rod DBC function using flash intensities that begin below the cone threshold and extend into the mesopic range, where both rods and cones contribute to the response. We also measured the photopic, cone only, response by saturating the rod response prior to delivery of high intensity flashes. From these data, we extracted the ERG b-wave amplitudes for rod-isolated (-3.6 log cd s/m^2^) and cone-isolated (1.4 log cd s/m^2^) responses for control and treated animals (Fig. 8). Example rod-isolated waveforms of *Lrit3^-/-^* animals treated with the RHO driven LRIT3 constructs (Fig. 8A) show only the full-length LRIT3 and ΔFN3 construct restored a response. The complete scotopic flash series is shown (Fig. 8B) and the rod-isolated amplitudes for all treated eyes (Fig. 8C) show that only Lrit3 and the ΔFN3 construct restored a response. To examine the function of the LRIT3 domains on cone photoreceptor function, we quantified the photopic ERG b-wave. Examples of photopic responses in a single eye treated with each construct are shown (Fig. 8D), as well as the results from the full photopic flash series for all eyes (Fig. 8B). These data show that the cone DBC signalplex is only functional when retinas are treated with either the full-length LRIT3 or the ΔFN3 construct.

**Figure 8.**
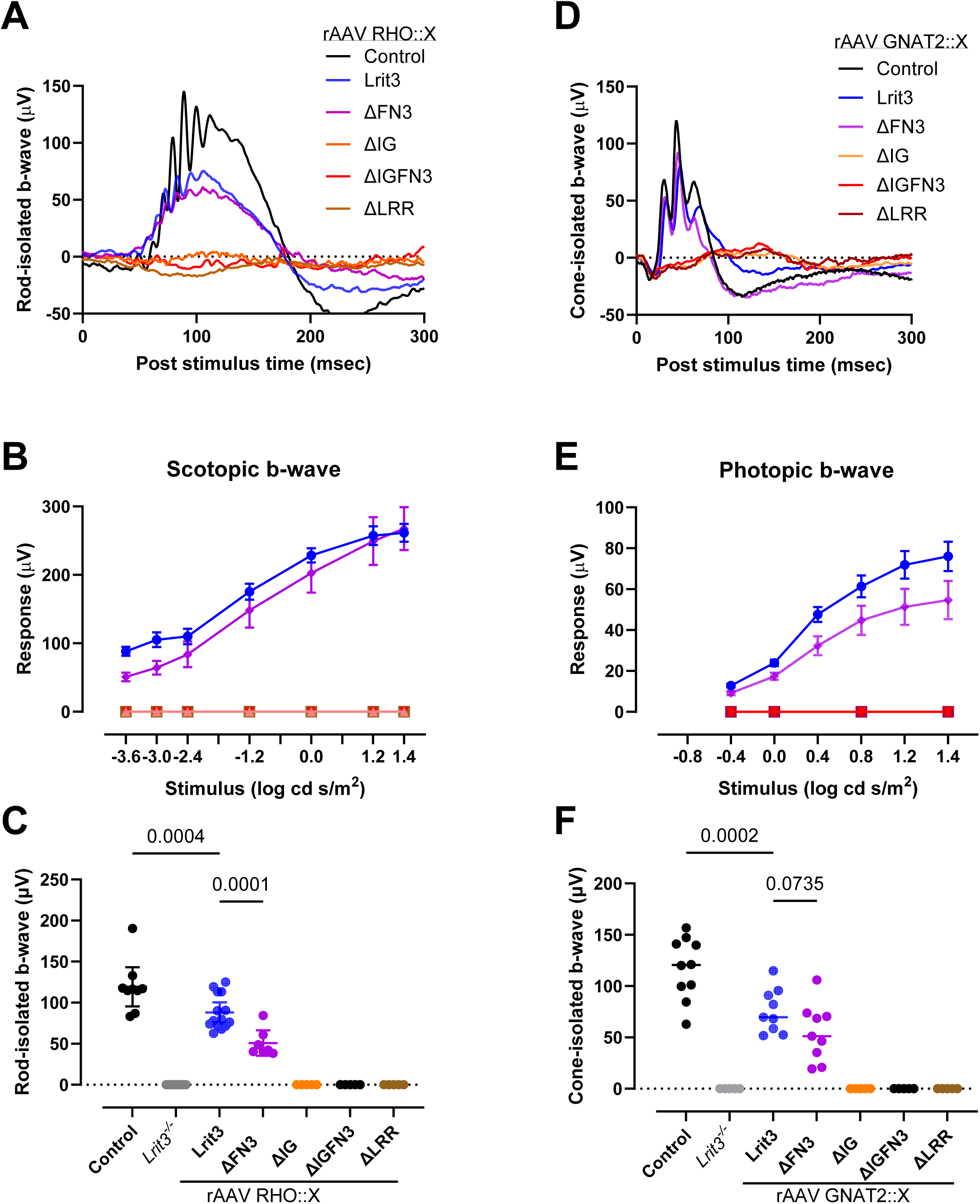
LRR, IG, and FN3 domains of LRIT3 are critical for synaptic transmission and retinal function. (A) Representative rod-isolated scotopic full-field electroretinogram (ffERG) waveforms recorded at -3.6 log cd s/m² from control mice (Wildtype treated with rAAV8 GRK1::GFP, n=9), *Lrit3*^-/-^ mice (n=8), and *Lrit3*^-/-^ mice treated with rAAV8 RHO-driven constructs: full-length Lrit3 (n=11), ΔFN3 (n=7), ΔIG (n=7), ΔIGFN3 (n=7), and ΔLRR (n=7). (B) Quantitative analysis of scotopic ffERG b-wave amplitudes across multiple flash intensities for experimental groups described in panel A. (C) Rod-isolated b-wave amplitudes from scotopic ffERG recordings at -3.6 log cd s/m² comparing all treatment groups from panel A. (D) Representative cone-isolated photopic ffERG waveforms recorded at 1.4 log cd s/m² from control mice (Wildtype treated with rAAV8 GRK1::GFP, n=9), *Lrit3*^-/-^ mice (n=8), and *Lrit3*^-/-^ mice treated with rAAV8 GNAT2-driven constructs: full-length Lrit3 (n=9), ΔFN3 (n=9), ΔIG (n=9), ΔIGFN3 (n=9), and ΔLRR (n=9). (E) Quantitative analysis of photopic ffERG b-wave amplitudes across multiple flash intensities for experimental groups described in panel D. (F) Cone-isolated b-wave amplitudes from photopic ffERG recordings at 1.4 log cd s/m² comparing all treatment groups from panel D. Data are presented as mean ± SEM.

A summary of the IHC and functional data for all the constructs is presented in Table 1. These data show that the FN3 domain is not required for trafficking in photoreceptors or assembly of a functional signalplex on all DBCs. In both rods and cones, the IG domain is not required for trafficking of the protein to the photoreceptor synapse, as indicated by IHC, but is insufficient to restore a functional signalplex on DBCs. The IG domain is not required for Nyctalopin expression on DBC dendrites; however, it is necessary for the proper expression of TRPM1.

**Table 1.** Function of LRIT3 domains.

| Lost in | RODs |  |  | CONEs |  |  |
| --- | --- | --- | --- | --- | --- | --- |
| *Lrit3*<sup>-/-</sup> | <u>Nyctalopin, TRPM1</u> |  |  | <u>Nyctalopin, TRPM1, mGluR6, GPR179</u> |  |  |
| Construct | Traffic | Restored | Function | Traffic | Restored | Function |
| Control | Y | All | Y | Y | All present | Y |
| *Lrit3*<sup>-/-</sup> | N | Nyc, TRP | N | N | All lost | N |
| ΔFN3 | Y | Nyc, TRP | Y | Y | Nyc, M6, TRP, GPR | Y |
| ΔIG | Y | Nyc | N | Y | Nyc, M6, GPR | N |
| ΔIGFN3 | Y | Nyc | N | Y | Nyc, M6, GPR | N |
| ΔLRR | Y | None | N | N | na | na |

\*Nyc=Nyctalopin, M6=mGluR6, TRP=TRPM1, GPR =GPR179

Finally, the LRR domain is essential for trafficking of LRIT3 to the synapse in cones, but not in rods.

## Discussion

We previously reported that LRIT3 is expressed presynaptically at both rod and cone terminals, where it trans-synaptically organizes the postsynaptic glutamate signaling complex on depolarizing bipolar cells (13, 17). The absence of LRIT3 leads to differential effects on the rod and cone DBC signalplexes that include mGluR6, GPR179, Nyctalopin, and TRPM1. We hypothesized that these differential effects may be mediated by the individual protein domains of LRIT3, which interact with its postsynaptic partners. In this study, we investigated the specific functions of LRIT3 extracellular domains, FN3, IG, and LRR. Frequently, such experiments are done in cultured cells because more manipulations are possible; however, the results in some cases do not reflect the *in vivo* situation. We demonstrated an example of this in a previous study, where *in vitro*, RG7 and RGS11 interacted with GPR179, whereas *in vivo*, only RGS7 showed this interaction (20). Therefore, in this study, we chose to use rAAVs to express various LRIT3 constructs *in vivo*. This allowed us to examine trafficking and functional outcome in the normal *in vivo* setting, namely, rods and cones.

LRIT3 is a single-pass transmembrane protein with three extracellular domains. The FN3 domain is most proximal to the membrane, followed by the IG and LRR domains. FN3 domains are protein-protein interaction domains that are widely distributed in mammalian proteins, particularly in cell adhesion molecules. LRIT3 mutants lacking the FN3 domain traffic and localize normally at both rod and cone terminals, and restore the DBC function as measured by the ERG b-wave. Included in the ΔFN3 construct is an unstructured 142 amino acid region (aa 341-482). Therefore, the FN3 domain and this linker region are not required for assembly or function of the postsynaptic DBC signaplex. The LRIT3 FN3/linker domain is highly conserved across all mammals, indicating that it plays a crucial role. One possibility is in positioning other proteins involved in glutamate dynamics that are expressed on photoreceptor terminals.

Evidence for this possibility is that in *Lrit3^-/-^* mice, there is reduced excitatory input to OFF bipolar cells, suggesting less glutamate reaches the flat synapses between cones and OFF BC dendrites (9). Further studies are needed to identify these potential interacting proteins.

The IG domain is located between the FN3 and LRR domains in LRIT3, and constructs lacking either the IG or both IG and FN3 domains have a similar impact on DBC signalplex assembly, namely, loss of TRPM1 from the signalplex. In the absence of the IG domain, Nyctalopin is localized correctly in both rods and cones, as are mGluR6 and GPR179 in cones. These data show that the LRIT3 LRR domain is required for mGluR6 localization to DBC dendrites, but not for its localization to rod BC dendrites. Instead, in rod BCs, the localization of mGluR6 is dependent on interactions with ELFN1 (10), mediated by glycosylation of the mGluR6 glutamate binding domain (21, 22). The trans-synaptic organization of mGluR6 and GPR179 in cone DBCs is complex and requires both LRIT3 and ELFN2 expression because in either ELFN1/2 or LRIT3 knockout mice, mGluR6 and GPR179 are absent (11, 15, 17, 23). Why the expression of ELFN1 and/or 2 in cones is insufficient to localize mGluR6 and GPR179, as it does in rod BCs, is unclear; however, this indicates that the mechanisms involved are different in rods and cones.

Previous studies reported that insertion of TRPM1 into the dendrites of DBCs, both rod and cone, required Nyctalopin (7), and a direct interaction between TRPM1 and Nyctalopin was reported (7, 24). However, we now show that by expressing the ΔIG construct, Nyctalopin is inserted into the DBC membranes, but this is insufficient to restore TRPM1 localization. Thus, Nyctalopin is necessary, but not sufficient, for normal positioning and function of TRPM1 in DBCs, which also requires the IG domain of LRIT3.

The LRR domain in LRR-containing proteins is involved in homomeric, heteromeric, or trans-synaptic protein-protein interactions (25). Interaction between nyctalopin and LRIT3 has been observed in a heterologous expression system (9). In the absence of the LRR domain, the mutant LRIT3 protein traffics normally to rod terminals. Still, it fails to restore nyctalopin and TRPM1 localization (Fig. 4). This may indicate that the LRR domain of LRIT3 interacts trans-synaptically with the LRR domain of nyctalopin. Alternatively, the LRR domain may be required to form homodimers, which then interact with their postsynaptic partner, nyctalopin, as is the case for ELFNs (26).

The trafficking of synaptic proteins to both pre- and postsynaptic locations is essential for proper function. This involves processing through the endoplasmic reticulum, then the Golgi, before being inserted into the membrane, where transsynaptic interactions occur, stabilizing the signaling complexes. These processes require specific domains, including the signal sequence, to ensure trafficking to the correct location. For LRIT3, the requirements for correct trafficking in rods and cones are different. In rods, only the intracellular domain is required, as shown by the trafficking of all constructs with the various extracellular domains deleted. In contrast, in cones, both the intracellular amino acids and the extracellular LRR domain are required. For ELFN1 and 2, their LRR domains interact to form homo- or hetero-dimers before they are inserted into the membrane, whereupon they interact with group III mGluRs (26). Whether this occurs for LRIT3 remains to be determined.

Our study is the first to elucidate the *in vivo* roles of LRIT3 domains in synaptic function and protein assembly in the retina. Our data reveal that each domain has a distinct function at both rod and cone metabotropic synapses. The LRR domain is essential for the localization of nyctalopin and TRPM1 at rod synapses and is critical for LRIT3 localization at cone synapses. Our findings do not indicate any involvement of the FN3 domain; however, both the IG and LRR domains are vital for ON pathway signaling. Future studies will be essential to elucidate the role of the FN3 domain in OFF pathway signaling, and why in cones both LRIT3 and ELFN2 are required for postsynaptic DBC signplex assembly.

## Experimental procedures

### Animals

All procedures were conducted following the Society for Neuroscience policies on the use of animals in research and the University of Louisville Institutional Animal Care and Use Committee guidelines. Animals were housed in the University’s AALAC-approved facility under a 12-hour light/dark cycle. The phenotype of the *Lrit3^-/-^* mouse line used in this study has been previously described (9, 13, 17).

Both male and female *Lrit3^-/-^* and C57BL/6J (WT) mice were used throughout the study. Mice were anesthetized with a ketamine/xylazine solution (118/11 mg/kg, respectively) diluted in normal Ringer’s solution before subretinal injections and ERG recordings. For enucleation of the eyes, mice were euthanized by CO_2_ exposure in accordance with AVMA guidelines. Data from mice with gross retinal damage from the injection procedure were excluded from the analyses.

### Antibodies

Antibodies used to label rod and cone synaptic proteins in the retina have been previously described (9, 13, 17) and are listed in Table 2. All antibodies were used at a dilution of 1:1000 and were validated for specificity by testing sections from the respective knockout mouse retinas. A GFP antibody was used to detect Nyctalopin-EYFP (27).

**Table 2.** Antibodies used.

| **Antibodies** | **Source** | **Identifier** |
| --- | --- | --- |
| Guinea pig anti-LRIT3 | Hasan et al, 2020 | N/A |
| Goat anti-mGluR6 | Gregg et al, 2023 | N/A |
| Rabbit anti-mCAR | Sheryl Craft, Zhu et al, 2002 | N/A |
| Rabbit anti-TRPM1 | Hasan et al, 2020 | N/A |
| Chicken anti-GFP | Invitrogen | Cat# A10262 |
| Sheep anti-GPR179 | Peachey et al. 2012 | N/A |
| Donkey anti-sheep IgG-AlexaFluor 488 | Life Technologies | Cat# A111015 |
| Donkey anti-rabbit IgG-AlexaFluor 647 | Life Technologies | Cat# A31573 |
| Donkey anti-guinea pig IgG-Cy3 | Millipore | Cat# AP193C |
| Donkey anti-rabbit IgG-AlexaFluor 488 | Life Technologies | Cat# A21206 |
| Donkey anti-goat IgG-AlexaFluor 647 | Life Technologies | Cat# A32849 |
| Donkey anti-chicken IgG-AlexaFluor 488 | Jackson Immuno Research Laboratories | Cat# 703-545-155;RRID: AB_2340375 |

### Retina preparation for immunohistochemistry

Retina sections and wholemounts were prepared following the methods described previously (13, 17). Briefly, mice were euthanized by CO_2_ inhalation followed by cervical dislocation. Eyes were enucleated, and the cornea and lens were removed. Retinas were dissected in PBS (pH 7.4) and fixed for 15-30 minutes in 4% paraformaldehyde diluted in phosphate buffer (0.1M PB). After fixation, retinas were washed three times in PBS for 10 minutes each, cryoprotected in a graded series of sucrose solutions (5%, 10%, 15% in 0.1M PB) for one hour at room temperature and then overnight at 4°C in 20% sucrose in 0.1M PB. Retinas were incubated in a mixture of OCT and 20% sucrose (2:1) for one hour prior to embedding. Cryoprotected retinas were frozen, and 18μm transverse sections cut using a cryostat (Leica Biosystems, Buffalo Grove, IL). Sections were mounted on Superfrost Plus slides (Thermo Fisher Scientific, Waltham, MA) and stored at -80°C. Immunohistochemistry methods have been previously described (9, 28). Retina sections and wholemounts were imaged using an FV-3000 Confocal Microscope (Olympus). Contrast and brightness adjustments were made using Fluoview Software (Olympus, Waltham, MA) or Photoshop (Adobe Systems, San Jose, CA).

### rAAV Production and Injection

To identify and annotate domains of LRIT3, we used the online SMART application (Simple Modular Architecture Research Tool, (29) and AlphaFold2 (30). Deletion mutants were cloned into the pcDNA3.1 vector. To express wild-type (WT) and deletion mutant LRIT3 proteins (ΔFN3, ΔIG, ΔIGFN3, and ΔLRR) in rods and cones, we employed recombinant adeno-associated viruses (rAAV8s) with gene expression controlled by the RHO and GNAT2 promoters, respectively (13, 17). The LRIT3 expression constructs were packaged into the rAAV8 capsid by VectorBuilder (Chicago, IL). The rAAVs expressing WT or deletion mutants of LRIT3 under the RHO promoter are designated as rAAV RHO::Lrit3, rAAV RHO::ΔFN3, rAAV RHO::ΔIG, rAAV RHO::ΔIGFN3, and rAAV RHO::LRR. For cone expression constructs, the RHO promoter was replaced by the GNAT2 promoter (19).

rAAV injections were performed by introducing 1 μl of the rAAV solution(1 x 10^13^ vg/ml in PBS) into the subretinal space of adult mice (postnatal day 35) using a specialized syringe (www.borghuisinstruments.com). Synaptic protein expression and retinal function were assessed four to eight weeks after rAAV injection.

### Electroretinography

Electroretinogram (ERG) recordings were performed as previously described (9, 31). Mice were dark-adapted overnight to ensure optimal sensitivity. They were then anesthetized with a ketamine/xylazine solution (118/11 mg/kg, respectively) diluted in Ringer’s solution, and prepared for recordings under dim red light to minimize additional light exposure. To prepare the mice, their pupils were dilated using topical applications of 0.625% phenylephrine hydrochloride and 0.25% tropicamide. The corneal surface was anesthetized with 1% proparacaine HCl. A feedback-controlled electric heating pad (TC1000; CWE Inc.) was used to maintain body temperature throughout the procedure. A contact lens with a gold electrode (LKC Technologies Inc.) was placed on the cornea and moistened with artificial tears (Tears Again; OCuSOFT, Gaithersburg, MD) to ensure proper electrical contact. Ground and reference needle electrodes were positioned in the tail and on the midline of the forehead, respectively. For the ERG measurements, scotopic responses were recorded with test flashes ranging from -3.6 to 1.4 log cd s/m², presented to dark-adapted mice. Photopic responses were measured with test flashes ranging from -0.8 to 1.4 log cd s/m² after a 5 minute light adaptation to a rod-saturating background of 20 cd/m².

### Rod and cone counts

The percentage of rod and cone photoreceptors expressing either WT or mutant LRIT3 in rAAV-treated *Lrit3^-/-^* retinas was quantified from wholemount immunostained images, as previously described (13). To identify all cone photoreceptors, retinas were stained with cone arrestin, while mGluR6 was used to label all rod synapses. LRIT3 staining was employed to mark transfected cells. For each rAAV expression construct, images were captured from three transfected retinas of three different mice using an FV-3000 confocal microscope with a 40x objective (NA = 1.4). Five images were taken per retina: one from each of the four quadrants and one from the central retina. These images were analyzed to determine the number of rods and cones expressing LRIT3 constructs. The total number of rod and cone synapses was estimated by counting mGluR6 puncta and cone arrestin, respectively. ImageJ software was used to count the puncta from maximum projections of z-stack whole-mount images.

### Statistical Analysis

Statistical analysis was performed using Prism 10.5.0 (GraphPad Software, Inc., La Jolla, CA). Details of the statistical methods are provided in the text and figure legends. Tukey post-hoc tests were applied to adjust for multiple comparisons where appropriate, and the adjusted p-values (p_adj_) are reported. Statistical significance was defined as p_adj_ ≤ 0.05.

## Data availability

All data are available upon reasonable request.

## Acknowledgements

Thanks to Timothy Hoffman for technical assistance.

## Author Contributions

Conceptualization, R.G.G. and N.H.; Investigation, N.H.; Writing, N.H. and R.G.G.; Funding Acquisition, R.G.G. and N.H.; Supervision, R.G.G.

